# Crb3 stabilizes activated Ezrin-Radixin-Moesin to organize the apical domain of multiciliated cells

**DOI:** 10.1101/2023.01.24.525309

**Authors:** Céline Burcklé, Juliette Raitière, Laurent Kodjabachian, André Le Bivic

## Abstract

Cell shape changes mainly rely on the remodeling of the actin cytoskeleton. Multiciliated cells (MCCs) of the mucociliary epidermis of *Xenopus laevis* embryos, as they mature, dramatically reshape their apical domain to grow cilia, in coordination with the underlying actin cytoskeleton. Crumbs (Crb) proteins are multifaceted transmembrane apical polarity proteins known to recruit actin linkers and promote apical membrane growth. Here, we identify the homeolog Crb3.L as an important player for apical domain morphogenesis in differentiating *Xenopus* MCCs. We found that Crb3.L is initially present in cytoplasmic vesicles in the vicinity of ascending centrioles/basal bodies (BBs), then at the expanding apical membrane concomitantly with BB docking, and finally in the ciliary shaft of growing and mature cilia. Using morpholino-mediated knockdown, we show that Crb3.L-depleted MCCs display a complex phenotype associating reduction in the apical surface, disorganization of the apical actin meshwork, centriole/BB migration defects, as well as abnormal ciliary tuft formation. Based on prior studies, we hypothesized that Crb3.L could regulate <u>E</u>zrin-<u>R</u>adixin <u>M</u>oesin (ERM) protein subcellular localization in MCCs. Strikingly, we observed that endogenous phospho-activated ERM (pERM) is recruited to the growing apical domain of inserting MCCs, in a Crb3.L-dependent manner. Our data suggest that Crb3.L recruits and/or stabilizes activated pERM at the emerging apical membrane to allow coordinated actin-dependent expansion of the apical membrane in MCCs.

## Introduction

Epithelia are lining the external surface and body cavities in animals and ensure exchanges with the extracellular environment as well as protection of internal organs (Buckley and St Johnston, 2022; Rodriguez-Boulan and Macara, 2014). This dual function relies on the organization of epithelial sheets. Epithelial cells are highly polarized, exhibiting structurally and functionally distinct apical and basolateral domains (Román-Fernández and Bryant, 2016). In a majority of epithelia in vertebrates, the apical surface forms microtubule-based protrusions called cilia (Apodaca, 2018). Primary cilia are antenna-like organelles perceiving a variety of environmental cues, including crucial signaling morphogens (Anvarian et al., 2019; Bangs and Anderson, 2017). In the ciliated epithelia lining the airways, the brain ventricles, or the reproductive tracts, the apical domain of multiciliated cells (MCCs) is covered by numerous cilia beating coordinately to propel biological fluids (Boutin and Kodjabachian, 2019; Spassky and Meunier, 2017). Thus, structural and/or functional alteration of cilia lead to rare diseases with pleiotropic symptoms called ciliopathies (Reiter and Leroux, 2017).

*Xenopus* is a powerful model for studying motile cilia biology (Walentek, 2021; Werner and Mitchell, 2012). The embryos are indeed enwrapped in a protective mucociliary epidermis composed of MCCs, bearing hundreds of motile cilia, alternating in a salt and pepper pattern with secretory cell types (goblet cells, ionocytes and small secretory cells (SSCs)) (Boutin and Kodjabachian, 2019; Walentek, 2021). The development of the *Xenopus* epidermis is a gradual process starting at segmentation/blastula stages with the partitioning of the prospective non-neural ectoderm into an outer epithelial layer made of goblet cells, and an inner mesenchymal layer where MCCs, ionocytes, SSCs and p63+ basal cells will be born (Deblandre et al., 1999; Dubaissi et al., 2014; Dubaissi and Papalopulu, 2011; Haas et al., 2019; Quigley et al., 2011; Stubb et al., 2006; Walentek et al., 2014). From mid-neurula onwards (Stages 14-31), intense morphogenetic events reshape the tissue, with the sequential insertion of MCCs, ionocytes and SSCs in the outer layer, where they complete differentiation (Haas et al., 2019). Successful intercalation of MCCs is powered by cell shape changes and forces generated by the actin and microtubule cytoskeletons (Boisvieux-Ulrich et al., 1987; Chuyen et al., 2021; Collins et al., 2021, 2020; Lemullois et al., 1988; Sedzinski et al., 2016; Werner et al., 2011). Prior to their intercalation, MCCs undergo random planar migration, constrained by homotypic repulsion, such that they eventually disperse and intercalate at regular intervals (Chuyen et al., 2021). MCC movement is confined to the grooves formed between overlying epithelial cells (Chuyen et al 2021). Radial intercalation involves the emission by MCCs of filopodia that pull on outer-layer junctions to probe their stiffness (Ventura et al., 2022). MCCs preferentially intercalate into stiff high fold vertices, where they provoke remodeling of the junction in preparation for apical expansion (Ventura et al., 2022). Once inserted into the outer layer, MCCs expand their apical domain, essentially via the 2D pressure exerted by a highly dynamic medial actin network generated by Formin 1 (Sedzinski et al., 2017, 2016).

While inserting into the outer layer, MCCs multiply centrioles deep in the cytoplasm, which are then encased via CP 110 and the ciliary adhesion complex (FAK, Paxillin, Vincullin, Talin) in an internal actin network that drives their apical migration (Antoniades et al., 2014; Walentek et al., 2016). Once docked at the apical surface, centrioles are named basal bodies (BBs). The dynamic ascension of centrioles relies on actomyosin contractility and is regulated by actors of the PCP signaling pathway, the core protein Dishevelled, the ciliogenesis and planar polarity effector (CPLANE) protein Inturned and the PCP effector RhoA (Adler and Wallingford, 2017; Boisvieux-Ulrich et al., 1990; Park et al., 2008, 2006). BBs serve as scaffold to initiate ciliogenesis, and are equipped with a basal foot and striated rootlets, which interact with actin filaments and microtubules to establish a regular lattice able to sustain intense ciliary beating strokes (Herawati et al., 2016; Mahuzier et al., 2018; Spassky and Meunier, 2017; Werner et al., 2011). In particular, an extensive reconstruction of the cortical actin meshwork in two layers enables the planar polarized anchoring of BBs to permit directional beating (Boisvieux-Ulrich et al., 1990; Ioannou et al., 2013; Mahuzier et al., 2018; Mitchell et al., 2007; Park et al., 2008, 2006; Werner et al., 2011). At stage 25, the most apical actin layer reaches its final density, actin bundles progressively encircle each BB, and from stage 26 onwards eventually mature in actin-based protrusion called microridges (Werner et al., 2011; Yasunaga et al., 2022). At stage 29-30, the definitive sub-apical actin layer connects each BB to its immediate posterior neighbour via ciliary adhesion complexes and to microridges via Ezrin (Antoniades et al., 2014; Werner et al., 2011; Yasunaga et al., 2022). This sophisticated actin architecture allows coordinated cilia beating, thus ensuring efficient flow production to clear the surface of the embryo from surrounding microbes (Dubaissi et al., 2018; Nommick et al., 2022).

To allow cell shape changes, deformation of the actin cytocortex must be coupled to deformation of the plasma membrane (Clark et al., 2014). The plasma membrane is attached to the underlying cytoskeleton via specific regulated linker proteins such as the ERM protein Ezrin (Clark et al., 2014; Pelaseyed and Bretscher, 2018). In *Xenopus* MCCs, overexpressed versions of Ezrin have been shown to accumulate in the newly expanding apical membrane of intercalating cells, and from there to be superimposed on the apical actin meshwork maturing into microridges (Yasunaga et al., 2022). Ezrin depletion in *Xenopus* MCCs induces a functional alteration of the actin cytoskeleton, impacting centriole/BB migration, apical microridge formation, and anchoring of BBs to microridges (Epting et al., 2015; Yasunaga et al., 2022). Thus, the fine-tuned localization of Ezrin correlates with the control of MCC terminal differentiation. However, the molecular mechanisms underlying the proper control of Ezrin subcellular localization in *Xenopus* MCCs are elusive.

The Crumbs (Crb) polarity proteins have pivotal functions in processes involving quick remodeling of the actin cytoskeleton in coordination with the adjacent membrane and are ERM biochemical and genetical interactors (Aguilar-Aragon et al., 2020; Bajur et al., 2019; Flores-Benitez and Knust, 2015; Gao et al., 2016; Kerman et al., 2008; Letizia et al., 2011; Médina et al., 2002; Salis et al., 2017a; Schottenfeld-Roames et al., 2014; Sherrard and Fehon, 2015; Simões et al., 2022; Tilston-Lünel et al., 2016; Vernale et al., 2021; Wei et al., 2015; Whiteman et al., 2014a). Biochemical studies have demonstrated that the Crb cytoplasmic tail possesses a FERM (Four-point one, ezrin, radixin, moesin) domain allowing ERM binding (Médina et al., 2002; Sherrard and Fehon, 2015; Tilston-Lünel et al., 2016; Wei et al., 2015; Whiteman et al., 2014a). Functional studies essentially performed in *Drosophila* embryos showed that Crb stabilizes the ERM protein Moesin at specific sub-apical domains depending on the nature of the remodeling tissues (Letizia et al., 2011; Salis et al., 2017a; Sherrard and Fehon, 2015). Crb and ERM both independently organize the apical cytocortex and interact genetically for dampening the actomyosin contractility during dorsal closure in *Drosophila* embryos (Bazellières et al., 2018; Fehon et al., 2010; Flores-Benitez and Knust, 2015). In vertebrates, the Crumbs family member Crb3 displays a quasi-ubiquitous epithelial expression (Bazellieres et al., 2009). Strikingly, Crb3- and Ezrin-deficient mice display the same abnormal intestinal phenotype, with villi fusion and microvilli atrophy (Charrier et al., 2015a; Saotome et al., 2004; Whiteman et al., 2014a). Both Ezrin and Crb3 have been shown to be required for ciliogenesis (Bazellières et al., 2018; Epting et al., 2015; Fan et al., 2004a, 2007; Hazime and Malicki, 2017; Omori and Malicki, 2006). However, how Crb3 and ERM proteins might cooperate to organize the apical domain and build protrusions, such as cilia and microvilli, in a vertebrate organism remains an open question.

We therefore took advantage of the mucociliary epidermis of the *Xenopus laevis* embryo to investigate Crb3 function in MCC terminal differentiation, and its interaction with ERM in establishing apical actin networks.

We uncovered that the two *crb3* homeologs, *crb3.L* and *crb3.S*, are expressed in an evolutive and exclusive pattern in the *Xenopus* mucociliary epidermis. Crb3.L is highly enriched in MCCs, where it localizes sub-apically in the vicinity of the ascending centrioles, at the apical plasma membrane, and in cilia. In contrast, Crb3.S is not detectable in MCCs, and is enriched in goblet cells and SSCs. Depletion of Crb3.L led to abnormal BB docking, correlating with a substantial alteration of the apical actin cytoskeleton, and consequently to defective ciliary tuft formation. We show that Crb3.L is required for phospho-activated ERM accumulation during the emergence of the apical MCC surface. Altogether, our data advocate for the requirement of Crb-ERM cooperation in fast actin remodeling-dependent processes such as centriole migration and apical surface emergence in MCCs.

## Materials and methods

### Cloning of crb3.L and crb3.S, constructs, mRNA, morpholinos®

Predictive ORFs of *crb3.L* and *crb3.S* of *Xenopus laevis* were identified by comparing genomic, RNA and protein sequences of *Xenopus tropicalis* and *Xenopus laevis*, as well as using the annotated versions of the genomes (Xenbase). cDNAs were prepared via reverse transcription (SuperSriptIII first strand synthesis, Invitrogen) from RNA extracted with silica membrane column purification (PureLink RNA mini kit Thermofisher).

PCR fragments were purified and cloned into pGEM-T Easy+, and subcloned in pCS2+ using the EcoR I cloning site to obtain the 29bp-crb3.L-pCS2+ vector, 29 bp of the 5’UTR were present in this first construct. The HA tagged crb3.L rescue construct resistant to 5’UTR morpholino, (HA-crb3.L-pCS2+), was constructed as follows. First, the HA epitope sequence was inserted immediately 3’ to the signal peptide sequence by site directed mutagenesis using the QuickchangeXL site directed Mutagenesis kit (Agilent) to obtain the following intermediate vector: 29bp-HA-crb3.L-pCS2+. To this end the following primers were used:

forward primer HAmutcrb3.L:

5’-CCTGTCATTAGTGAAAGCTTACCCATACGATGTTCCAGATTACGCTCAGAATGTCACCACTTCAG-3’

reverse primer HAmutcrb3.L:

5’-CTGAAGTGGTGACATTCTGAGCGTAATCTGGAACATCGTATGGGTAAGCTTTCACTAATGACAGG-3’

The insertion of an optimal Kozak sequence with simultaneous deletion of the 5’UTR remnant were performed by PCR using as a template the restriction fragment obtained by the digestion of 29bp-HA-crb3.L-pCS2+ by EcoRI.

The following primers were used.

forward 5’ GAGAGAATTCCGCCACCATGCTCGATTATCTA

reverse primer 5’GAGACTCGAGACTAGTGATTTGTAAGGCGC.

PCR fragments were digested, purified and cloned into PCS2+ using the EcorI-XhoI cloning sites.

Accuracy of the sequences were verified by sequencing.

The plasmids encoding tracers: PCS2+-RFP-caax, PCS2+-GFP-GPI were from lab stocks. PCS107-GFP-CP110 was a kind gift of Peter Walentek (Center for biological system analysis Universitätsklinikum Freiburg, Freiburg, Germany).

Linearized plasmids were used as template for *in vitro* synthesis of mRNA using the SP6 m message machine kit (Ambion) and purified using the Megaclear Kit (Life technology Ambion).

Two independent morpholinos® were designed targeting either the 5’UTR of crb3.L, named *5’UTR-crb3.L-mo*, encoded by 5’-ACAGTAACAGGGTAGGGACGCA-3’ or the translational initiation codon named *ATG-crb3.L-mo* encoded by 5’-TTAGTAGATAATCGAGCATGTGGAC-3’.

### *Xenopus* embryo injections

Wild-type or albino *Xenopus laevis* females were obtained from Biological Resources Center (CRB France or NASCO, USA). Eggs were fertilized *in vitro*, and embryos were de-jellied and reared as previously described (Nommick et al., 2022). For microinjections, embryos were placed in a 5% Ficoll buffered solution (Hatte et al., 2018; Nommick et al., 2022). To specifically target the epidermis, 8-cells embryos were injected in a ventral blastomere of the animal pole. For depletion experiments, embryos were injected with 15 or 30 ng of *5’UTR-crb3.L-mo* or *ATG-crb3.L-mo* respectively, together with 200 pg of *gfp-gpi* mRNA. For rescue experiments, *5’UTR-crb3.L-mo* was co-injected with 200 pg of *mrfp-caax* mRNA and the HA tagged resistant *crb3.L* mRNA.

### Immunohistochemistry

Embryos were staged and arrested at appropriated stages according to Faber and Nieuwkoop (Faber and Nieuwkoop, 2020). For stage 21-22 embryos were incubated at 13°c until late afternoon the day of injection and then placed at 16°c for 2 days. For stage 28, embryos were incubated at 13°c for 4 days. For embryos that did not hatch, the vitelline membrane was manually removed with tweezers before proceeding to fixation. Fixation conditions varied according to the antibodies and/or the probes (see tables A, B, C). After fixation, embryos were immunostained *in toto*, or further processed for cryosections. Cryopreservation was achieved by incubation in serial graded sucrose solutions (5 to 30%). Embryos were then embedded in Optimal Cut Temperature medium (OCT VWR chemicals), snap frozen in a dry ice ethanol bath, and stored at −80°C, 5 to 10 μm thick cryosections were obtained with cryostat.

**Table A:** primary antibodies.

| Primary antibodies | Specie/isotype | reference | Dilution | fixation |
| --- | --- | --- | --- | --- |
| Ezrin (phospho T567) | Rabbit polyclonal | # 47293 abcam | 1/200 | PFA 4% 1h RT |
| γ-Tubulin | mouse IgG1 | ref T5326 Sigma-Aldrich | 1/800 |  |
| Crb3.S | rabbit polyclonal | This work | 1/200 | MEMFA 2h RT followed by 100% methanol at -20°C |
| Crb3.L | rabbit polyclonal | This work | 1/200 |  |
| Centrin | mouse IgG2 | ref 04-1624-Merck-Millipore | 1/1000t | PFA 4% 1h RT<br>Or<br>MEMFA 2h RT followed by 100% methanol at -20°C |
| α-acetylated Tubulin | mouse IgG2b | T6793 Sigma-Aldrich | 1/500 |  |
| RFP | rat IgG2a | 5F8 Chromotek | 1/500 |  |
| GFP | chicken polyclonal | GFP-1020 AvesLab | 1/500 |  |
| HA | mouse IgG1 | # 49969 Abcam | 1/500 |  |

**Table B:** secondary antibodies.

| Secondary antibodies alexa fluor | reference | dilution |
| --- | --- | --- |
| anti mouse IgG-488 | A-11029 Life technologies | 1/500 |
| anti mouse IgG- 546 | A-11003 Life technologies |  |
| anti mouse IgG- 647 | A-21235 Life technologies |  |
| anti mouse IgG1- 647 | A-21240 Life technologies |  |
| anti mouse IgG2a-647 | A-21241 Life technologies |  |
| anti-mouse IgG2a-488 | A-21131 Life technologies |  |
| anti mouse IgG2b-647 | A-21242 Life technologies |  |
| anti rat IgG(H+L)-568 | A-11077 Life technologies |  |
| anti rabbit IgG-488 | A-11008 Life technologies |  |
| anti rabbit IgG-546 | A-11035Life technologies |  |
| anti chicken IgY-488 | A11039 Life technologies |  |

**Table C:**
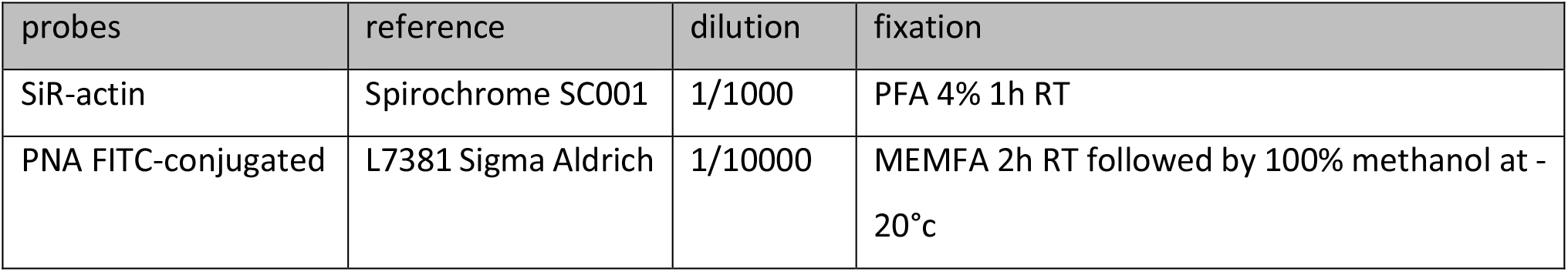
fluorescent probes.

| probes | reference | dilution | fixation |
| --- | --- | --- | --- |
| SiR-actin | Spirochrome SC001 | 1/1000 | PFA 4% 1h RT |
| PNA FITC-conjugated | L7381 Sigma Aldrich | 1/10000 | MEMFA 2h RT followed by 100% methanol at -20°C |

For whole mount immunostaining, buffers and detergents varied with the antibodies or probes. For staining including p-ERM and SiR actin, embryos were washed in a tris buffer saline (TBS) NP-40 0,05% solution, and blocked in TBS-NP40 0,05% with 15% fetal calf serum (FCS). For the other staining, embryos were washed in maleic acid and triton 0,1% (MABX), and blocked in MABX with 15% FCS. Next, embryos were usually incubated overnight at 4°C with the first antibodies (see table A). Embryos were then extensively washed for 4 to 6 hours, incubated 30 minutes to 1 hour in blocking buffer, then transferred in the solution containing the appropriate combination of goat secondary alexa-fluor coupled antibodies at 1/500 for 1h30-2 hours (see table 2). After incubation, embryos were shortly washed and mounted in antifading medium (mowiol-DABCO) between slide and coverslip. Exceptions to this protocol were the increased incubation time for anti-Crb3 antibodies (3-4 days, 4°C), addition of SiR-actin 1/1000 to the fixative solution (1 hours) and with the secondary antibodies (1h30 hours) (see table C).

For Crb3.L immunostaining on cryosections, reagents and conditions were the same as described above. Peanut agglutinin (PNA) was incubated on the slide for 15 minutes after the secondary antibody incubation and wash (see table C).

### Generation of Crb3 homeolog L and S specific antibodies

Crb3 homeolog L and S specific antibodies were obtained by immunizing rabbits using the speedy protocol from Eurogentec (Seraing, Belgium). The synthetic peptides identical to the ectodomain of Crb3.L (H-QNVTTSAPDRLSESAR-C) or Crb3.S (H-QNVTTPAPGKLSESA-C) were coupled with the KLH (keyhole limpet hemocyanin) carrier protein on their C-terminal part. These peptides were used for affinity purification of the immunized rabbits. Validation of the antibodies was demonstrated via immunizing peptide competition assays for Crb3.L and Crb3.S and depletion through morpholino knockdown for Crb3.L (see supplementary data).

Competition assays were performed using the three following conditions. The antibodies against Crb.3 were incubated overnight at 4 °C with either the immunizing peptide, or an unrelated peptide (QTISDPGEEDPPVSKC present in *Oopsacas minuta* type IV collagen, negative control) or only MABX buffer (positive control). The molar ratio between antibodies and peptides was 1 mol for 25 mol, respectively, and antibodies were used at 1:200 (2.6 μg/mL). After centrifugation (21,000 g, 4 h, 4 °C) to get rid of potential antibody-peptide complexes, supernatants were used to perform the immunostaining protocol. Incubation of antibodies with the immunizing peptide caused the loss of immunostaining unlike incubation with *Oopsacas minuta* type IV collagen.

### Imaging and quantification

Confocal images were acquired using ZEISS LSM 510 and 780, Leica SP5 and SP8 confocal microscopes with 63 x oil objectives. Sequential or simultaneous laser excitations were applied depending on fluorophores, combinations, spectra, staining brightness and persistence.

### Basal Bodies

#### Apically anchored BBs

Centrin staining was used for BBs detection. Apical surfaces of individual MCCs marked by the tracer corresponding to regions of interest (ROI) were segmented manually. Maximum intensity projection of the 3 planes framing the MCCs apical surface was applied to Centrin pictures and converted to 8-bits format. Automated detection of Centrin dots was performed in the ROI using the function find maxima>point selection of Fiji. For this purpose, images were acquired with 63 x oil objective applying a 2,5 zoom and z slice interval of 0,4 μm.

#### Non apically anchored BBs

Z series were set to frame the totality of the Centrin staining in MCCs. The number of Z-slices containing Centrin dots was used as a proxy for the apico-basal dispersion of BBs in the cytoplasm. These values were converted in distances from the cell apex. For this purpose, images were acquired with 63 x oil objective applying a 2,5 zoom and z slice interval of 0,4 μm.

#### p-ERM and F-actin signals

Fluorescence was measured on 8-bit pictures using the mean gray value function of Fiji on cells, from which surfaces were manually delineated, on stacks combining the three most apical cell plans. p-ERM signal quantification was performed in mosaic tracer-injected control and crb3.L-depleted embryos to circumvent non-biological interindividual variations of the staining intensity. Measures of mean pixel intensity of pERM staining per cell were performed on confocal plan including at least two tracer positive (cti) and negative MCCs (ctni) in control embryos, as well as two 5’UTR-crb3L-mo + tracer positive (moi) and negative (moni) MCCs in morphant embryos. Then, the average intensity (mean pixel intensity) of each cell population was calculated for each plan. The absence of variation of the average intensity between cti and ctni was a prerequisite for interpreting a variation between moi and moni.

F-actin staining and imaging conditions were very reproducible allowing comparison of MCCs F-actin fluorescence between control (tracer only) and morpholino (5’UTR-crb3L-mo + tracer) injected embryos.

For p-ERM and F-actin stainings, images were acquired with 63 x oil objective and z slice interval of 0,5 μm.

#### Transmission electron microscopy

Stage 28 Embryos were processed for electron microscopy as previously described (Revinski et al., 2018). 80 nm sections were made with a Leica Ultracut UC7 (Leica, Germany). Images were acquired using a Tecnai G2 (Thermofisher, USA) microscope and a Veleta camera (Olympus Japan).

#### Statistical analysis

Statistical analyses were performed with RStudio (version 1.4.1717). Before comparing the mean of variables, normality and homoscedasticity were evaluated with Shapiro and Barlett tests, respectively. When data followed normal and homoscedastic distribution, Student’s t-test was applied for comparing two groups. When data did not complete these two conditions, non-parametric tests were used. For two-groups comparison, Wilcoxon test was used. For comparison including more than two groups, Kruskal-Wallis tests were used, followed by post hoc Wilcoxon-test with Bonferroni correction to examine differences between two means. Distribution are depicted as Box-plot, box represent the interquartile range (50% of the distribution) and whiskers highlight 1,5 interquartile range, median is shown. Chi-squared test was applied for percentage comparison. For all tests, significance threshold was set at p<0,05. * for p<0,05, ** for p<0,01, *** for p<0,005, **** for p<0,001

## Results

### Crb3.S and Crb3.L display distinct and dynamic expression pattern in the mucociliary epidermis of *Xenopus laevis* embryos

In *Xenopus laevis*, there are two *crb3* homeologs, *crb3.S* and *crb3.L*, with highly conserved DNA (85 % identity) and protein (85 % identity) sequences. As no tools were available to observe the precise endogenous localization of Crb3 proteins in *Xenopus laevis*, we generated and validated homeolog specific antibodies. To do so, we used the extracellular domain of Crb3.S (QNVTT<u>P</u>AP<u>GK</u>LSESA) and Crb3.L (QNVTT<u>S</u>AP<u>DR</u>LSESA<u>R</u>) as immunogens, as this region is the most divergent between the two proteins (non-identical amino acids are underlined). To assess the specificity of the Crb3.L and Crb3.S antibodies, we performed competition assays with the corresponding immunogenic peptides, which caused the extinction of immunofluorescent (IF) signals in whole-embryos (supplementary figures 1A and 2). By combining these new tools with Utrophin labelling that stains the actin meshwork, acetylated-Tubulin that stains stable cytoplasmic microtubules and cilia, and Centrin that stain centrioles and BBs, we were able to show that Crb3.L and Crb3.S have dynamic and exclusive pattern in the mucociliary epidermis of *Xenopus* embryos (figures 1 and 2).

**Figure 1:**
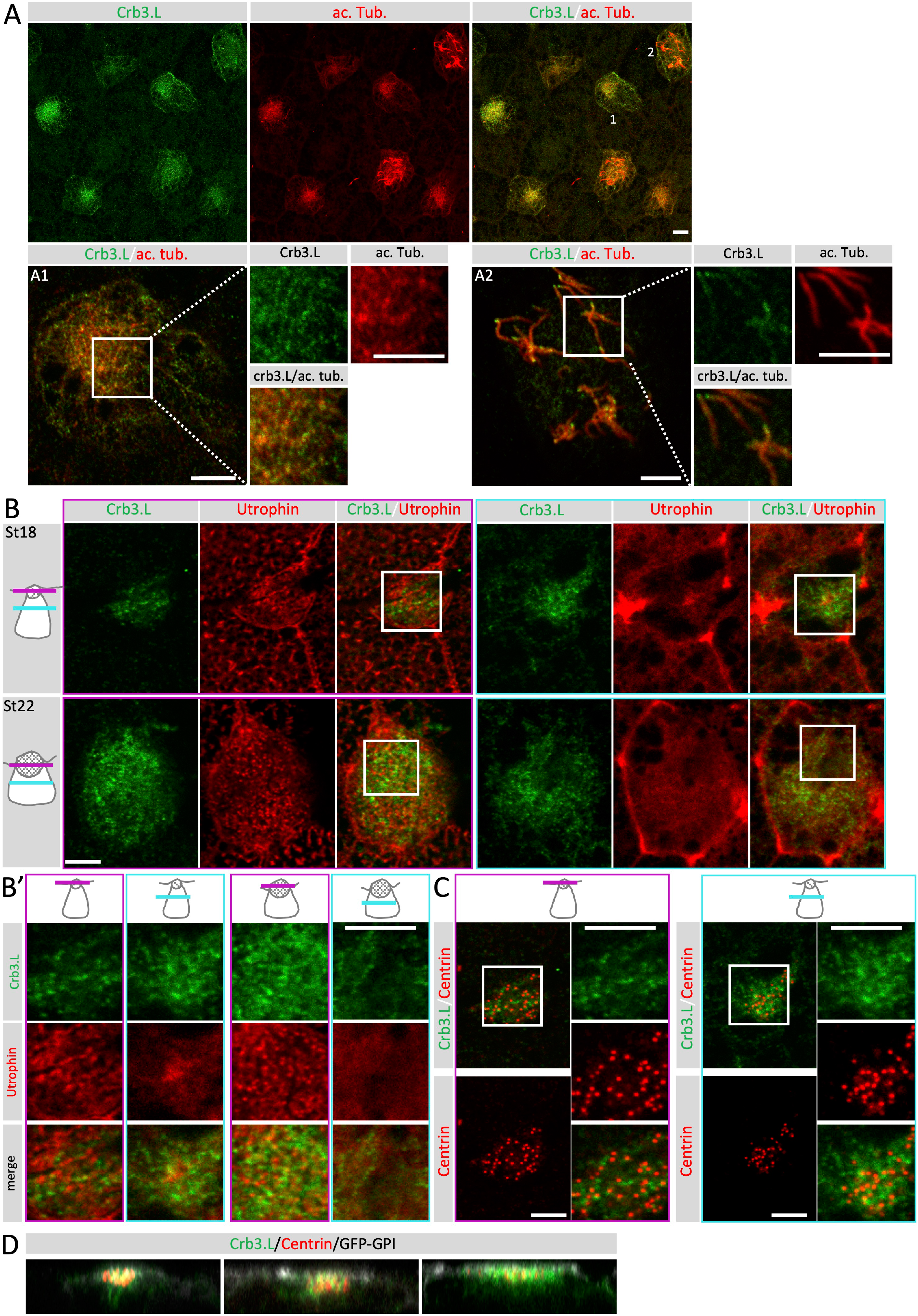
Crb3.L is expressed in MCCs expanding their apical surface and during ciliogenesis. **A-D: Micrographs of MCCs during insertion, expansion of the apical domain and early ciliogenesis in stage 18 embryos.** Crb3.L is labelled with our custom made anti-Crb3.L. Scale bars are 10 μm. **(A): Crb3.L vesicles are in close association with microtubular tracks in MCCs** Maximal intensity projection of en face view of whole-mount of St 18 *Xenopus laevis* embryos at low magnification. Stable microtubules are labelled with anti-acetylated a-tubulin antibodies (ac. Tub.) A1, A2: Single confocal section at higher magnification of the MCC 1 and 2 pointed in A. A1 section at the level of the tubulin network in cell 1 sub-apical domain. A2 section at the level of cilia in cell 2. Inserts on the right-side panel A1 and A2 are cropped magnified micrographs of the region of interest (ROI), corresponding to white squares drawn in A1 and A2. Inserts display separate channels and overlay for better appreciation of the localization of Crb3.L regarding stable microtubules. Note the dotty staining of Crb3.L following microtubule tracks in the cytoplasm (A1) and the ciliary axoneme (A2). **(B): Crb3.L vesicles localize within the remodeling actin meshworks in MCCs** F-actin is labelled via injection of *utrophin-gfp* mRNA (Utrophin). (B): upper row, single x, y confocal sections of an emerging MCC (apical surface 63μm^2^) in a stage 18 embryo. Purple framed panels are at the level of the newly formed apical domain as depicted on the scheme (purple line indicates the level of the section). Cyan frame panels are at the level of the internal actin meshwork as depicted on the scheme (cyan line indicates the level of the section). (B) lower row, single x, y confocal sections of a an emerged MCC (apical surface 178 μm^2^) in a stage 22 embryo. Purple framed panels are at the level of the maturing apical actin meshwork as depicted on the scheme (purple line indicates the level of the section). Cyan frame panels are at the level of the internal actin meshwork as depicted on the scheme (cyan line indicates the level of the section). (B’) cropped magnified micrographs of the ROI, corresponding to white squares drawn in B; cells and levels of section are indicated on the scheme heading each column. **(C): Crb3.L vesicles localize in the vicinity of the ascending centrioles** Centrioles and BBs are labelled by anti-Centrin antibodies. Single x, y confocal sections of the St18 MCC shown in B. Purple framed panels are at the level of the centrioles/BBs that have reached the apical domain as depicted on the scheme (purple line indicates the level of the section). Cyan frame panels are at the level of the internal centrioles/BBs as depicted on the scheme (cyan line indicates the level of the section). **D: Crb3.L localization changes are synchronous to centriole/BB ascension and apical surface emergence in MCCs.** x-z optical sections through a series of MCCs with expanding apical surface. Micrographs are ranked from the smallest (left) to the largest apical cell surface. Note the progressive shift of Crb3.L from an internal position when centrioles/BBs are deep in the cytoplasm to the apical domain when BBs are docked at the apical surface.

**Figure 2:**
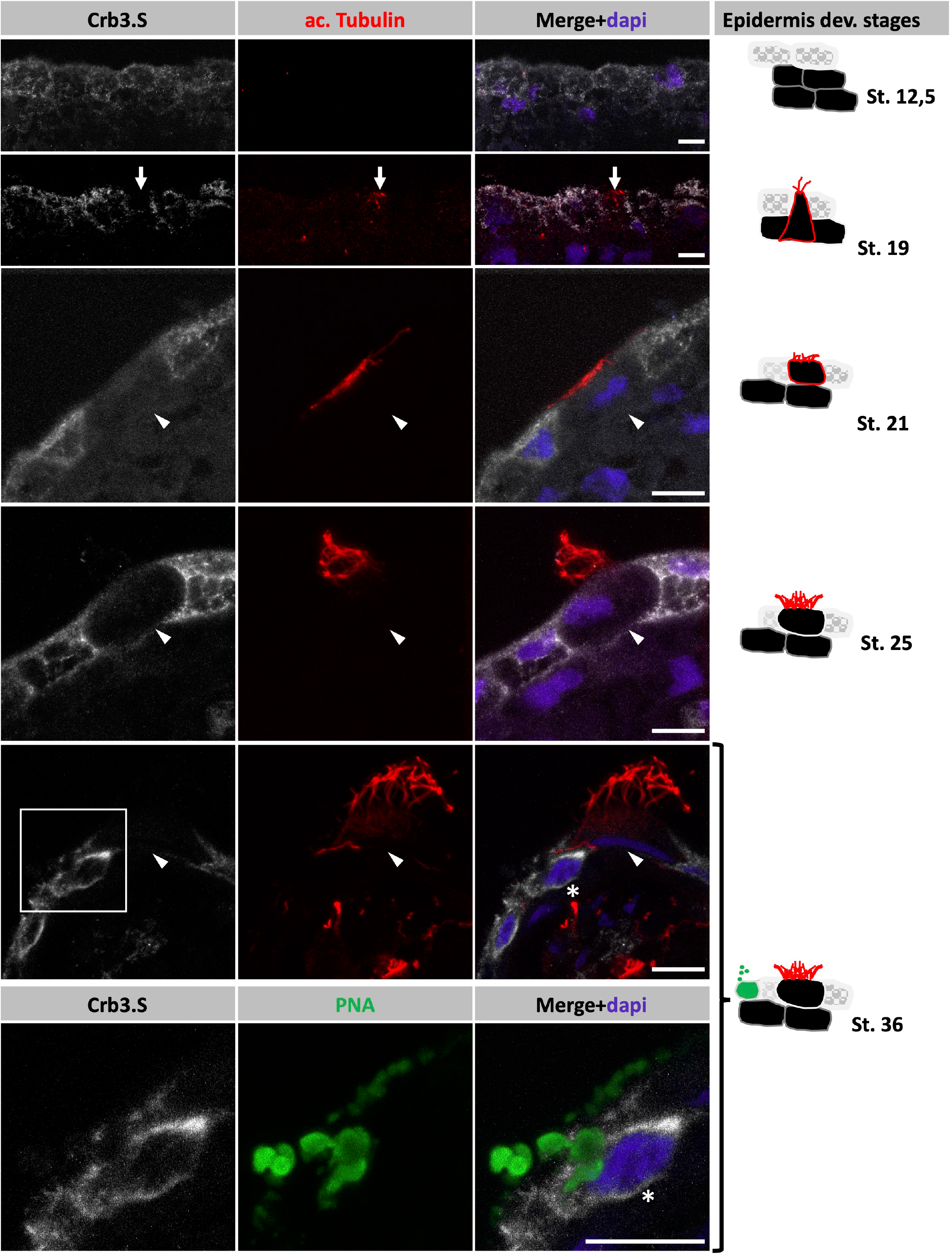
Crb3.S has an early, dynamic, and wide expression during morphogenesis of the mucociliary epidermis. A: Frozen sections displaying Crb3.S custom-made antibody staining during morphogenesis of the mucociliary epidermis. Schemes on the right depict *Xenopus* epidermis morphogenetic changes according to the developmental stages. Nuclear staining is DAPI, cilia are labelled with anti-acetylated a-Tubulin (ac-Tubulin), peanut agglutin (PNA) stains small and large secretory vesicles of goblet cells and SSCs, respectively. (white arrows) newly formed cilia, (white arrowheads): MCCs, (star): SSC. Scale bars are 10 μm.

We focused on the expression of Crb3.L in intercalating MCCs (figure 1). In inserting MCCs, Crb3.L-positive vesicles start to accumulate deep in the cytoplasm close to microtubule arrays (figure 1 A, A1), at the level of the internal actin meshwork (figure 1 B, B’, D), in the vicinity the ascending centrioles (figure 1 C,D). Next, as apical emergence and conjoint centriole ascension proceeds, Crb3.L is progressively redistributed to the expanding apical domain proximate to actin filaments (figure 1, B, B’, C, D) and appears in the ciliary shaft of growing cilia (figure 1, A2).

In contrast, Crb3.S exhibits an earlier and wider expression, being detected in all cells of the outer layer during gastrulation (figure 2 A). From gastrulation onwards, Crb3.S expression level is prominent in goblet cells, albeit at variable levels, presumably absent from ionocytes and particularly high in SSCs (figure 2, supplementary figure 2B). Strikingly, it is very low or absent from intercalating and mature MCCs (figure 2, supplementary figure 2).

Based on this analysis and the central function of CRB proteins in cell shape changes empowered by quick actin remodeling, we decided to evaluate the role of Crb3.L in the morphogenesis of the apical domain of MCCs.

### Crb3.L is required for proper ciliogenesis

To address the function of *crb3.L* during ciliogenesis, we used an antisense morpholino knockdown strategy with two morpholinos targeting either the translation initiation site (*ATG-crb3.L-mo*) or the 5’untranslated region (*5’UTR-crb3.L-mo*) of the *crb3.L* mRNA. The efficiency of *5’UTR-crb3.L-mo* was evidenced by the extinction of Crb.3L IF staining in whole-embryos, in both MCCs and non-MCCs (supplementary figure 1B). To unravel a potential effect on ciliogenesis, we examined stage 28 embryos, when MCCs are mature, with a well-developed ciliary tuft (Fig 3, control). Crb3.L depletion strongly affected the aspect of the ciliary tuft in MCCs, with a reduction in the number of cilia. Quantitative analysis revealed that ciliogenesis was significantly impaired upon injection of either of the *crb3.L* targeting morpholinos, with 30% (confidence interval 24-37%, *5’UTR-crb3L-mo*) to 39 % (confidence interval 26-53%, *ATG-crb3L-mo*) of MCCs forming abnormal ciliary tufts with a reduced number of cilia (figure 3). As the two *crb3.L* targeting morpholinos displayed comparable effects, *5’UTR-crb3.L-mo* was used in the rest of the study.

**Figure 3:**
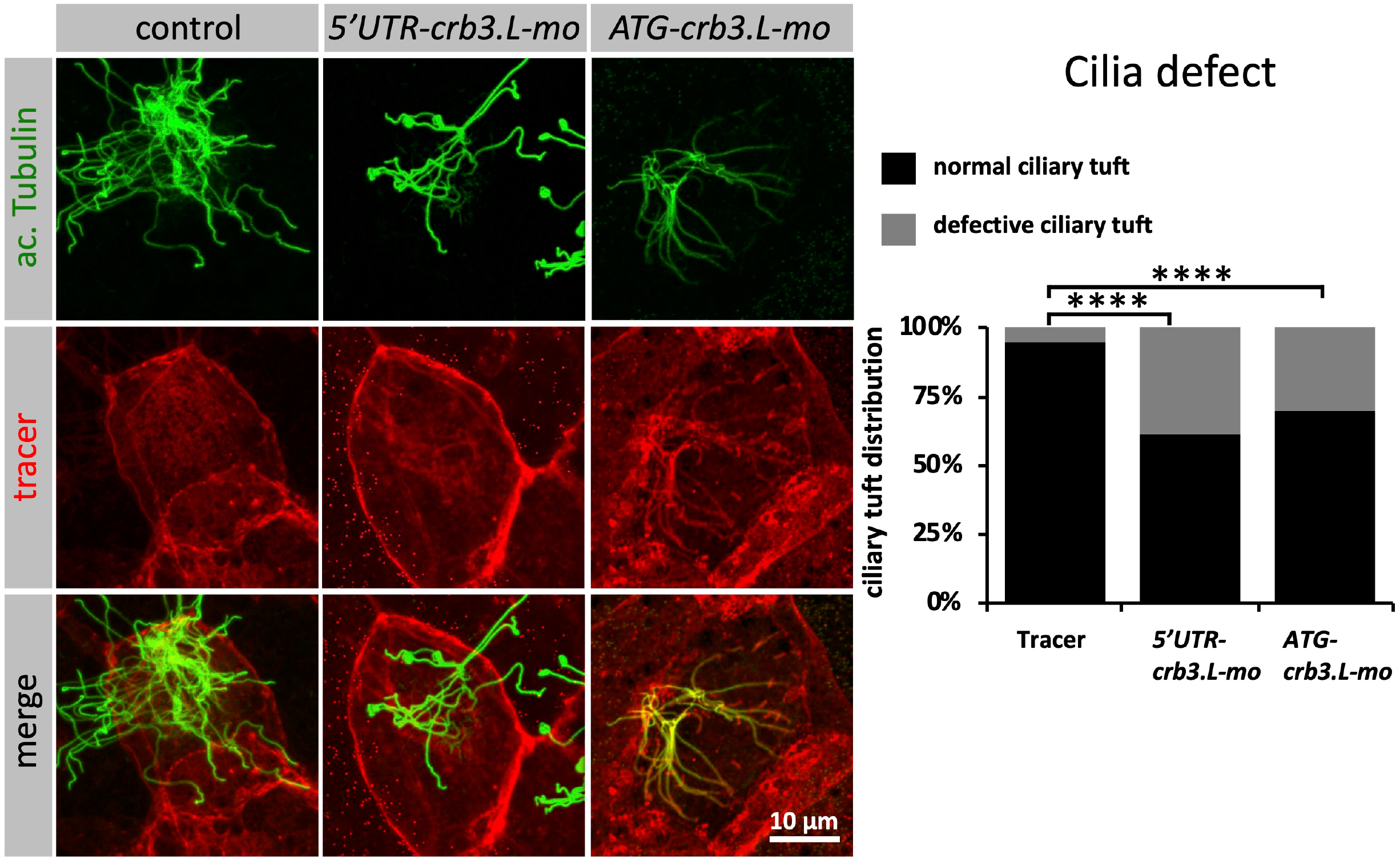
Crb3.L is required for ciliogenesis in MCCs. A: En face view of stage 28 embryos, control embryos are injected with 200 pg of *m-RFP* mRNA, morphants are co-injected with 200 pg of *m-RFP* mRNA and either 15 ng of 5’UTR targeting morpholino (*5’UTR-crb3.L-mo*) or 30 ng of ATG targeting morpholino (*ATG-crb3.L-mo*) and *mRFP* mRNA. Cilia are labeled with anti-acetylated a-Tubulin. B: Bar charts showing the quantification of the cilia defects. For quantification, the value 1 was assigned to well-furnished cilia tufts as in control (see anti-acetylated a-Tubulin control picture, the value 0 was assigned to defective cilia tuft displaying an obviously low cilia number such as in morphant (see anti acetylated a-Tubulin in morphants). χ^2^test, followed by pairwise comparison using Bonferroni correction. Controls (8 embryos, 94 MCCs), *5’UTR-crb3.L-mo* (8 embryos, 56 MCCs), *ATG-crb3.L-mo* (20 embryos; 225 MCCs). ****p<0,001. Pictures and bar charts from one representative experiment out of 3 experiments.

### Crb3.L is required for proper apical positioning of BBs

As correct BB apical docking is a prerequisite for proper ciliogenesis, we compared BB position in tracer-injected control and morphant MCCs. Two complementary approaches were used to observe the organization of BBs: immunolabelling and transmission electron microscopy (TEM). At stage 28, in the control situation, BBs cover most of the apical area, with quite a regular distribution (figure 4, A, B, E, F, I). In contrast, in *5’UTR-crb3L-mo*-injected MCCs, defaults in BB position and distribution are readily apparent in confocal and TEM images. BBs remained stuck into the cytoplasm (figure 4 C, D, G, H, I) or unevenly scattered, forming clumps when they reached the apical surface (figure 4 I). To quantify the penetrance of this phenotype, we analyzed two parameters: the number of BBs reaching the apical domain and the dispersion of BBs along the apico-basal axis (figure 4 J, K). The number of BBs reaching the apical surface was drastically decreased in morphant compared to control MCCs (figure 4 J). Furthermore, BBs were located significantly deeper into the cytoplasm of morphant compared to control MCCs (figure 4 K). Importantly, both aspects of the BB phenotype were significantly rescued by the injection of a HA-tagged Crb3.L construction insensitive to the *5’UTR-crb3.L* morpholino (figure 4 J, K).

**Figure 4:**
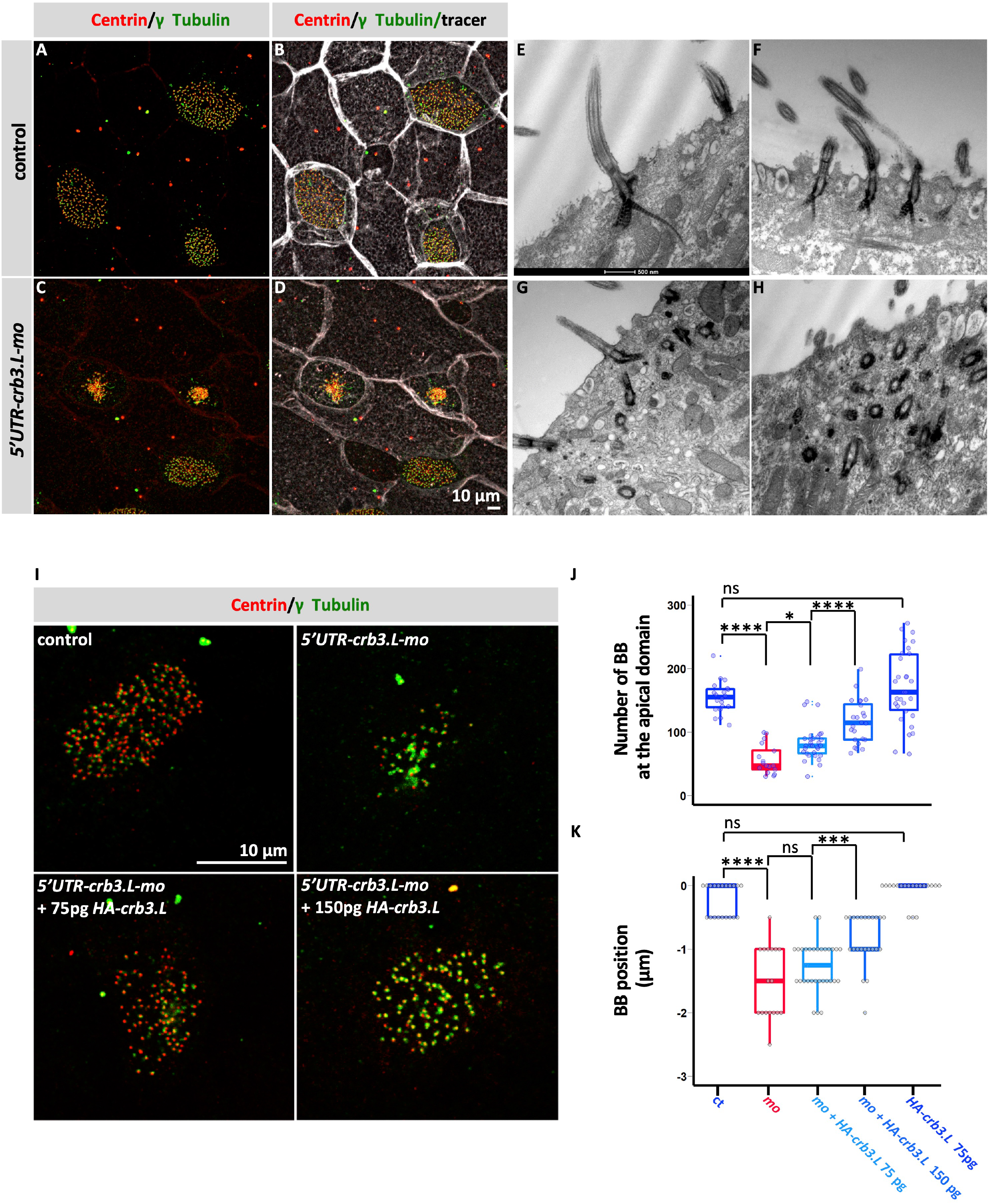
crb3.L is required for proper BB migration and docking. (A-D): Maximum intensity projection of en face view stage 28 embryos. BBs are labeled with anti-Centrin (centriole) and anti γ-Tubulin (basal foot) antibodies. Control embryos are injected with 200 pg of *GFP-GPI* mRNA, morphants are co-injected with 200 pg of *GFP-GPI* mRNA and 15 ng of *5’UTR-crb3.L-mo*. (E-H) TEM picture of the skin of stage 28 embryos. (E, F) control embryos, (G, H) morphant embryos. Note the even distribution of BBs covering most of the apical surface in control embryos (A, B, F, I) whereas in morphant embryos apically located BBs occupy a restricted part of the cell surface, are clumped together and stay deep in the cytoplasm (C, G, H) (I-J) Rescue experiment showing correction of the BB migration/docking defect with morpholino insensitive HA tagged *crb3.L mRNA*. (I) Most apical slice of a representative MCC, in the 4 following injection conditions: control (no injection), co injection of 15 ng *5’UTR-crb3.L-mo* and tracer, co injection of 15 ng *5’UTR-crb3.L-mo* and 75 pg *HA-crb3.L* mRNA and tracer, co injection of 15 ng *5’UTR-crb3.L-mo* and 150 pg *HA-crb3.L* mRNA and tracer. (J, K) To assess the rescue, we estimated two parameters: the number of BBs docked on the apical surface (upper graph J) and the position of the BBs relative to the cell apical surface (lower graph K). Injection conditions are common to the two graphs, and are depicted below the BBs position graph. mo refers to *5’UTR-crb3L-mo*. (J) The number of apically located BBs was automatically counted with the find maxima tool of the Fiji software. (K) To estimate BB position relative to the apical cell surface, the number of plans occupied by at least 3 BBs were counted. Uninjected control (n=20), co injection 15 ng of *5’UTR-crb3L-mo* and tracer (n=19), co injection of 15 ng *5’UTR-crb3L-mo* and 75 pg *HA-crb3.L* mRNA and tracer (n=24), co injection of 15 ng *5’UTR-crb3L-mo* and 150 pg *HA-crb3.L* mRNA and tracer (n=32), 75 pg *HA-crb3.L* mRNA (n=21), n= number of cells. Results are presented with Box-plots, box displays the interquartile range (50% of the distribution) and whiskers highlight 1,5 interquartile range, median is shown. *p<0,05, ***p<0,005, ****p<0,001

All these data demonstrated that Crb3.L is essential for an efficient apical migration and/or docking of centrioles/BBs.

### Crb3.L is required for proper cortical actin meshwork organization in mature MCCs

Migration/docking of BBs is a stepwise process relying on proper actin meshwork dynamics and organization. In several models, Crb has been shown to regulate actin dynamics (Bazellieres et al., 2009; Flores-Benitez and Knust, 2015; Röper, 2012a; Salis et al., 2017b; Sherrard and Fehon, 2015; Simões et al., 2022). As Crb3.L-positive vesicles repartition displayed dynamic changes correlating with actin meshwork remodeling (Figure 1), we hypothesized that Crb3.L could regulate actin cytoskeleton organization in MCCs. We therefore assessed cortical actin meshwork organization in stage 28 control and *5’UTR-crb3L-mo-injected* MCCs. In both uninjected (red contour arrowheads) and injected (orange contour arrowheads) control cells, the labeled actin meshwork was dense, structured, and anchored to the cell junctions (figure 5 A, B, I, J). In *5’UTR-crb3L-mo-injected* MCCs (filled orange arrowheads), the central actin meshwork appeared much dimer, fragmented and largely disconnected from the cell junctions (figure 5, E, F, K, L). Measurement of actin staining intensity confirmed a drastic decrease of about 50% in morphant compared to control MCCs (figure 5M). At this stage, the mean area of the apical domain of MCCs was not significantly different between the two groups, suggesting that in mature MCCs there is no strict correlation between the apical cell surface area and the actin meshwork density (figure 5N).

**Figure 5:**
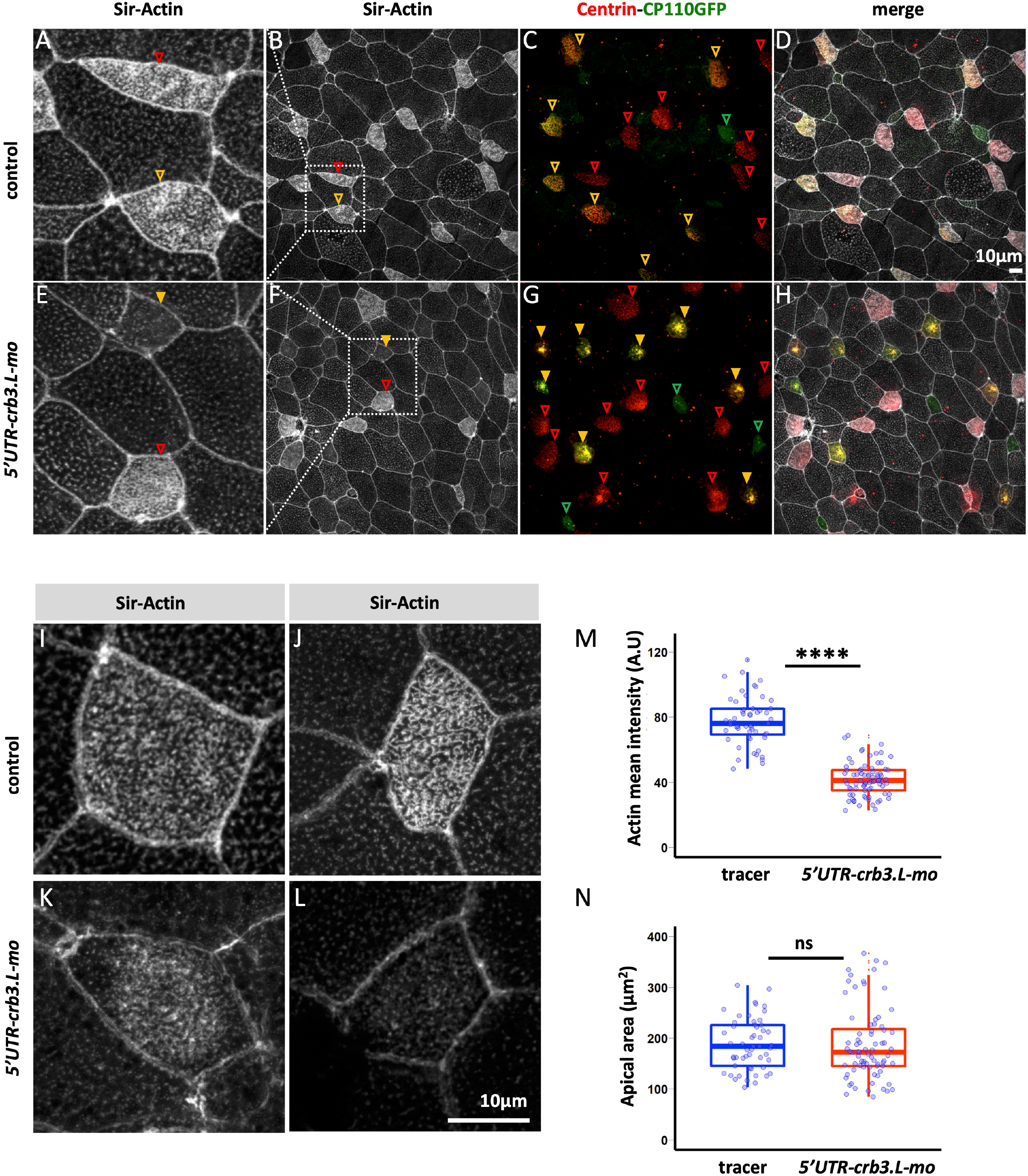
Crb3.L is required for proper organization of the actin cytoskeleton in MCCs. (A-H) Maximum intensity projection of whole-mount embryos, actin is stained with the Sir-actin probe, BBs are detected with anti-Centrin antibodies, *cp110-GFP* mRNA is a tracer labelling BBs in injected cells. (A-D) Mosaic control injected with *cp110-GFP* mRNA, (I-H) mosaic morphants co-injected with *5’UTR-crb3L-mo* and *cp110-GFP* mRNA. (C, G) Uninjected MCCs are detected via Centrin endogenous staining displayed in the red channel in both mosaic controls and morphants (red contoured arrowheads). Non-MCCs cells are labelled with Cp110-gfp displayed in the green channel (green contoured arrowheads). Injected MCCs are revealed by doubly labelled BBs (endogenous Centrin+ injected *cp110-GFP* tracer) and thus appear in a yellow to orange range of color. Orange contoured arrowheads point to control tracer injected MCCs. Orange filled arrowheads point to (*5’UTR-crb3L-mo* + *cp110-GFP* tracer) injected MCCs. BB intracytoplasmic aggregation results in a bright orange-yellow signal (orange filled arrow). (A-E) crop image of the ROI delineated by the white dashed line in B and F. (I-L) Higher magnification of the actin meshwork in control and morphant MCCs. (I) Quantification of the actin cytoskeleton defect, medial regions were manually delineated excluding the junctions, mean actin intensity was measured with Fiji and is presented in arbitrary unit (number of experiments >3). Results are presented with Box-plots, box displays the interquartile range (50% of the distribution) and whiskers highlight 1,5 interquartile range, median is shown. ****p<0,001

These results show that in mature MCCs, Crb3.L is required for proper organization of the cortical actin meshwork, but seems dispensable to reach the final apical surface area.

### Endogenous pERM associates dynamically with ascending centrioles/BBs and the apical actin meshwork

We and others have shown that ERM and CRB proteins cooperate for apical domain construction (Aguilar-Aragon et al., 2020; Bajur et al., 2019; Flores-Benitez and Knust, 2015; Letizia et al., 2011; Médina et al., 2002; Salis et al., 2017a; Sherrard and Fehon, 2015; Tilston-Lünel et al., 2016; Wei et al., 2015; Whiteman et al., 2014a). Previous studies in *Xenopus* have demonstrated that Ezrin-depleted MCCs exhibit a very similar phenotype to the one we observed in Crb3.L-depleted MCCs (Epting et al., 2015).

Thus, we hypothesized that Crb3.L might affect the expression and/or the subcellular localization of activated Ezrin. As a first step to evaluate this possibility, we analyzed the localization of phospho-activated ERM during MCC differentiation. In control MCCs, pERM expression was detectable in the protruding membrane starting its insertion into the outer layer (Figure 6 St18, supplementary figure 6). At this stage, pERM staining was also detected in close proximity to the ascending centrioles/BBs, (supplementary figure 6 stage 21). As apical expansion proceeded, pERM formed bright patches on the expanding apical surface (Figure 6 St22, 25), then the patches coalesced in a more structured network superimposed on the mature actin meshwork (Figure 6 St25-30). Accumulation of pERM at cell-cell junctions was observed only in fully mature cells (Figure 6 St 30).

**Figure 6:**
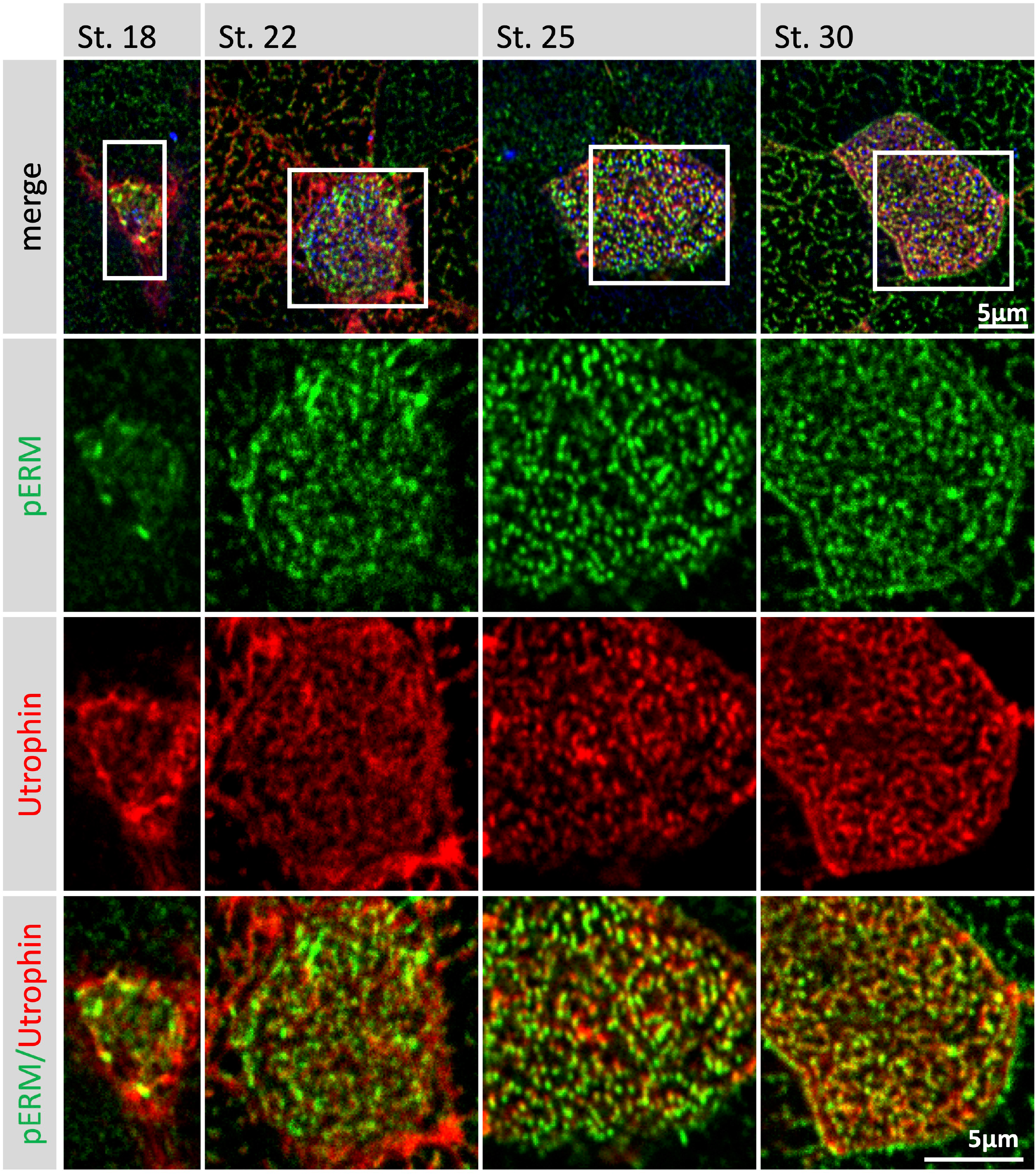
Endogenous pERM expression in MCCs. Single confocal section of whole-mount embryos between stages 18 and 30, focusing on the emerging and maturing apical domain of MCCs. pERM detection is coupled Utrophin-GFP to label actin. ROIs are shown by white squares in the first row, cropped magnified pictures corresponding to these ROIs are presented in the following rows.

### Crb3.L regulates the localization of pERM at the apical membrane

The similar apical distribution of Crb3L and pERM suggests that Crb3.L could control the localization of activated pERM to stabilize the nascent actin network during apical expansion. We thus compared the distribution of pERM in control and Crb3.L-depleted immature MCCs undergoing apical expansion. Crb3.L depletion induced a 33,2% (+/-11 sd) decrease of apical pERM signal intensity (figure 7 A, B, D). In correlation with this effect, Crb3.L depletion provoked a 30 % (+/- 10 sd) decrease in the average apical surface of newly emerged cells. These results are in contrast to the apparently preserved area of stage 28 morphant MCCs (figure 7 C, figure 5 N), and suggest that non-Crb3.L-dependent processes might rescue the initial defect in apical expansion.

**Figure 7:**
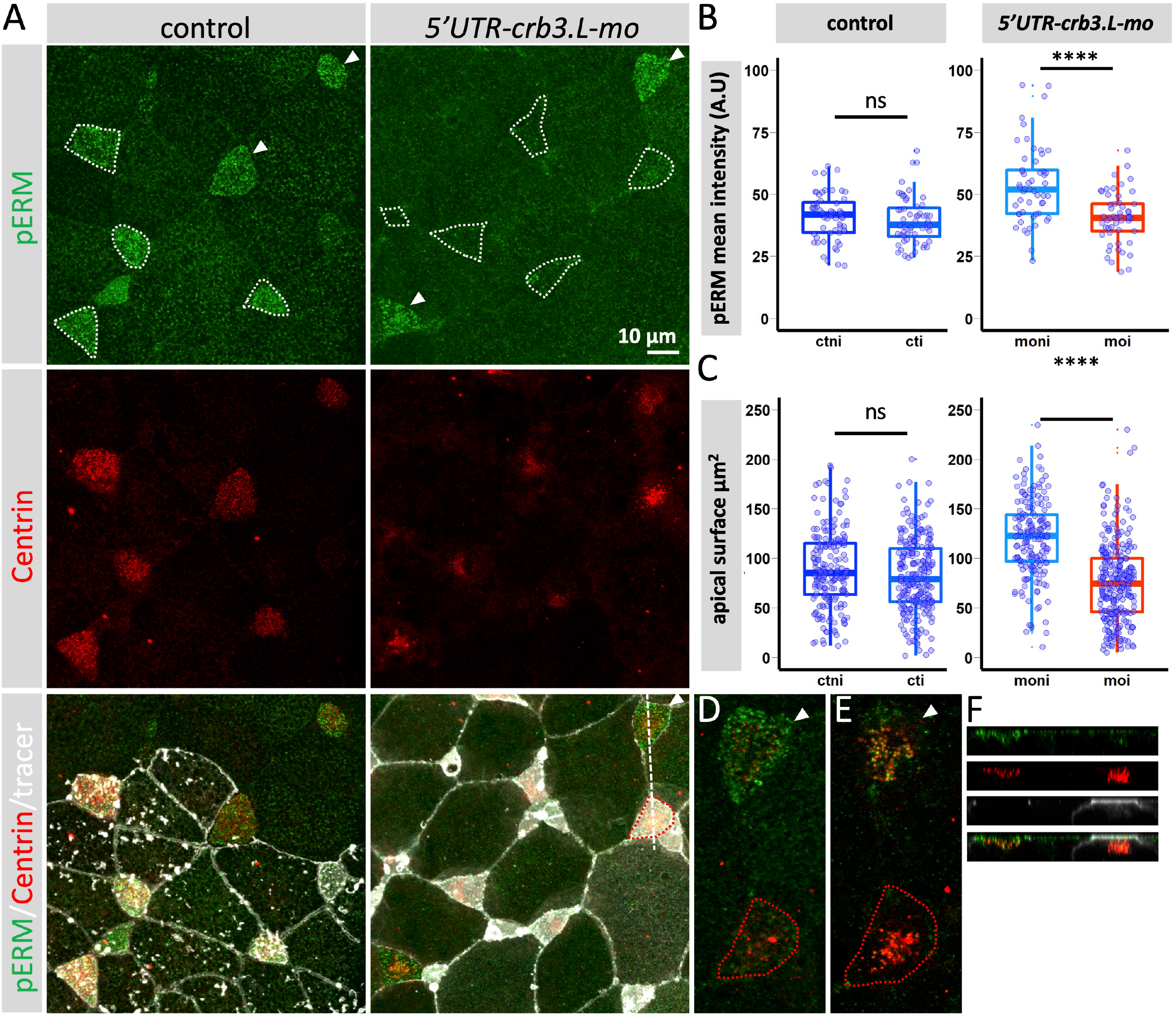
Crb3.L recruits pERM at the developing apical surface of MCCs and is important for apical surface expansion. (A) Maximum intensity projection of a whole-mount mosaic tracer injected control and *5’UTR-crb3.L* morphant stained with anti-pERM antibodies. BBs are detected with anti-Centrin antibodies, GFP-GPI is the tracer for injected cells in control and morphant stage 21 embryos. Tracer Injected, or tracer + morpholino injected MCCs are encircled by dotted lines. Uninjected MCCs are indicated by white arrowheads. (B) Quantification of pERM intensity in mosaic control and 5’UTR-crb3L morphants. To avoid inter- and intra-embryos non-biological staining variations, only mosaic injected regions were considered for quantification in control and *5’UTR-crb3.L* morphants at stage 21. Mosaic frames containing more than 2 injected and 2 uninjected MCCs were used for quantification to allow calculation of the average intensity of pERM staining in each cell population per frame. Absence of variation of the staining in mosaic controls was a prerequisite for considering a variation observed in mosaic morphant as significant. Box plots present the average intensity of pERM staining per frame in MCCs of mosaic controls (left) and mosaic morphants (right). uninjected control MCCs (**ctni**), tracer injected MCCs (**cti**) MCCs, uninjected morphant MCCs (**moni**), *5’UTR-crb3L-*morpholino + tracer injected MCCs (**moi**). Pictures and graphs from one representative experiment out of 3 experiments. Mosaic controls (n=9, 187 uninjected cells and 246 injected cells, 56 frames), mosaic morphants (n=9, 176 uninjected cells and 268 injected cells, 53 frames). (C) Quantification of the size of the apical domain in mosaic control and *5’UTR-crb3L* morphants at stage 21. Measures are performed on the pictures selected for figure 7B. Absence of variation in the apical surface in mosaic controls was a prerequisite for considering a variation observed in mosaic morphant as significant. Box plots present the distribution of the individual measures of apical surface in MCCs of mosaic controls (left) and mosaic morphants (right). uninjected control MCCs (**ctni**), tracer injected MCCs (**cti**) MCCs, uninjected morphant MCCs (**moni**), 5’UTR-crb3L-morpholino + tracer injected MCCs (**moi**). Pictures and graphs from one representative experiment out of 3 experiments. Mosaic controls (n=9, 187 uninjected cells and 246 injected cells, 56 frames), mosaic morphants (n=9, 176 uninjected cells and 268 injected cells, 53 frames). (D-F) pERM recruitement at BBs in a mosaic morphant embryo (D, E) higher magnification of single focal plane at the level of the apical surface (D), and inside the cells at the level of ascending BBs (E). The uninjected cell is pointed by the white arrowhead, the injected cell is surrounded by the red dotted line. (F):x,z optical section according to dotted line in A, channel color code is the same as in A. Note that this configuration (uninjected and injected cells with similar apical surface and ascending BBs) is an extremely rare event making difficult quantification of pERM at BBs.

These data advocate for Crb3.L being required for the stabilization of pERM on the emerging apical surface of MCCs, so as to bridge newly formed actin filaments to the apical plasma membrane, and promote early apical expansion.

## Discussion

In this study, thanks to newly homemade polyclonal antibodies, we show that the Crb3.L homeolog is highly expressed in MCCs, with a dynamic pattern of expression at the nexus of BBs and the intensively remodeling cytoskeleton of microtubules and actin. Crb3.L-depleted MCCs displayed a complex phenotype associating BB migration defects, reduction in the apical surface, disorganization of the apical actin meshwork, thus precluding normal ciliogenesis. We further unveil the transient association of endogenous activated ERM with ascending centrioles/BBs and its recruitment to the growing apical surface during the last step of MCCs intercalation. The loss of apical enrichment of pERM paralleled the initial reduction in apical domain expansion in Crb3.L-depleted MCCs. Our data advocate for Crb3.L stabilizing Ezrin in an open active conformation at the apical surface of MCCs, thus allowing coordinated growth of the apical membrane with extensive actin cytoskeleton remodeling.

### Crb3 homeolog expression in the *Xenopus* mucociliary epidermis

So far, studies addressing the question of Crb3 tissue and subcellular localization in *Xenopus laevis* relied on the expression of a C-terminally GFP-tagged version of *Xenopus tropicalis* Crb3. The amino acid sequence of *Xenopus tropicalis* Crb3 displays 83 and 81% identity with Crb3.L and Crb3.S respectively (Chalmers et al., 2005; Wang et al., 2013, p. 6). This construct allowed the description of the dynamic changes in Crb3 subcellular localization during differentiation and morphogenesis of the non-neural ectoderm, but did not address the specific question of its subcellular localization in MCCs. To determine which homeolog was predominantly expressed in MCCs, and to describe their subcellular localization, we generated antibodies targeting each one of the proteins and controlled their specificity by immunizing peptide competition assays and loss of function experiment for Crb3.L. These new antibodies allowed us to unveil the distinct but complementary expression of Crb3.L and Crb3.S in the developing mucociliary epidermis of *Xenopus laevis* embryos, supporting a division of labor scenario following duplication.

We confirm that a large pool of Crb3, namely Crb3.S, is indeed located in numerous small intracellular vesicles (Wang et al., 2013). Moreover, the progressive restriction of the endogenous Crb3.S to the outer layer at the time of its mucociliary differentiation is in accordance with the published pattern based on the expression of *tropicalis* Crb3-GFP (Wang et al., 2013). These data advocate that Crb3.S is a marker signing the beginning of the differentiation of the mucociliary epidermis. Later on, the expression of Crb3.S became very variable among the different cell types and surprisingly extremely high in the SSC.

To date, very few SSC markers have been described and they are either transcription factors such as Foxa1, or allow the detection of secreted compounds (Otogelin-like, HNK1, IPTKb, serotonin, Xpod) (Dubaissi et al., 2014; Kurrle et al., 2020; Walentek et al., 2014). Thus Crb3.S is, so far, the sole transmembrane protein described to be highly expressed in SSCs. It extends the short list of bona fide SSC markers, and offers the unique advantage to visualize cell shape via endogenous staining of vesicles and cytoplasmic membranes.

The SSCs are central to the epithelial anti-infective barrier function in *Xenopus laevis* embryos via secretion of anti-microbial substances (Dubaissi et al., 2014; Walentek et al., 2014). Therefore, it would be interesting to study the impact of Crb3.S depletion on the secretory function of the SCC as well as its functional consequences on innate immunity.

Until now, Crb3 localization in ciliated cells focused on its localization in the shaft of primary cilia (Fan et al., 2007, 2004b; Hazime and Malicki, 2017; Sfakianos et al., 2007). Accordingly, functional studies on Crb3 revealed its function in the regulation of intraciliary transport (Fan et al., 2004b; Hazime and Malicki, 2017; Sfakianos et al., 2007) and ciliary membrane composition (Hazime and Malicki, 2017). Here, we do observe a spotty staining of Crb3.L along the axoneme, suggesting that Crb3.L might also regulate ciliary transport/ciliary shaft composition in motile cilia of MCCs.

Before being detected in cilia, Crb3.L accumulates in the sub-apical domain of the intercalating MCCs, in the vicinity of the ascending centrioles/BBs, as well as the newly expanding apical membrane. These subcellular localization data are well in accordance with the compound phenotype we observed: the association of defective centriole/BB ascension, alteration of the apical actin meshwork and apical surface reduction. These data suggest that in MCCs, Crb3 is not exclusively required for ciliary transport, as it has been described for primary cilia (Fan et al., 2004b; Hazime and Malicki, 2017; Sfakianos et al., 2007). Our work points that Crb3.L is likely an iterative player of the stepwise process of multiciliogenesis, being subsequently required for centriole/BB ascension, BB anchoring/fusion to the apical membrane, apical actin meshwork construction, and finally cilia building/maintenance.

### Crb3.L vesicles might modulate ERM phosphorylation status at BBs

Accumulation of Crb3.L vesicles in the vicinity of centrioles/BBs at the time of their migration, and defective BB positioning in Crb3.L morphants, clearly suggest that Crb3.L regulates the BB ascension process. Our work provides some important hints to understand how Crb3.L and ERM might contribute to this process.

Here, we show that endogenous pERM staining is intense at ascending centrioles/BBs in inserting cells or cells with small apical surfaces. Remarkably, intense endogenous pERM staining at centrioles/BB is restricted in a short time window (stage 18-21) and does never label all the MCCs in a field, suggesting a transient and asynchronous recruitment of pERM at BB/centrioles. Thus, the endogenous pERM staining we observed is well in accordance with the model of dynamic and transient recruitment/activation of pERM at BBs allowing their ascending progression along actin filaments (Epting et al., 2015). In this model the Elmo-Dock-Rac1 module act as part of a molecular switch required for Ezrin dephosphorylation (Epting et al., 2015). The BB phenotype of Crb3.L-depleted cells is similar to what has been observed in Ezrin-depleted MCCs (Epting et al., 2015). Thus Crb3.L vesicles might regulate ERM phosphocycling at BBs in several ways. Via direct interactions, Crb3.L could stabilize ERM in their active phosphorylated conformation at BBs as described in *Drosophila*, or at the expanding apical surface of MCCs (this work, Flores-Benitez and Knust, 2015; Sherrard and Fehon, 2015). Indirectly, Crb3.L vesicles could function as signaling platforms bringing uncharacterized kinase(s) required for pERM activation at BBs, as described in enterocytes, where apical recycling endosomes carry the kinases that activate Ezrin at the plasma membrane (Dhekne et al., 2014; Gould and Lippincott-Schwartz, 2009). Moreover, Crb3.L might interfere with the Elmo-Dock-Rac1 module required for Ezrin dephosphorylation, as Crumbs can antagonize Rac1 activity during some developmental processes in *Drosophila* embryos (Chartier et al., 2011; Sollier et al., 2015). Understanding how Crb3.L vesicles modulate the molecular switch of Ezrin phosphorylation during centriole/BB ascension clearly needs further investigations.

### Crb3.L vesicles might coordinate apical membrane expansion with the growth of the apical actin cytocortex

A widely accepted function of Crumbs is to control the size of the apical domain in various model systems *in vivo*. In *Drosophila*, Crb loss-of-function leads to smaller apical surfaces of tracheal and pupal wing cells (Salis et al., 2017a; Skouloudaki et al., 2019). Crb might also selectively participate to the expansion of apical domain subdivision such as the stalk and rhabdomere regions of *Drosophila* photoreceptors, the inner segment of the photoreceptor and the apical microvilli-like protrusions of mechano-sensory neurons in zebrafish, and the microvilli of enterocytes in mice (Charrier et al., 2015a; Desban et al., 2019; Omori and Malicki, 2006; Pellikka et al., 2002; Richard et al., 2009; Whiteman et al., 2014a). In summary, Crb is required for apical growth in tissues undergoing intense morphogenetic events.

In *Xenopus laevis*, overexpression of Crb3 leads to expansion of the apical domain of epithelial cells in the early embryo (Chalmers et al., 2005). However, the question of Crb3 contribution to the very dynamic process of the MCC apical expansion has not been evaluated. Here, we show that prior to apical MCC expansion, Crb3.L labels vesicles accumulating in the subapical domain of inserting cells. Further on, Crb3.L relocates from this vesicular pool to the apical membrane at the time of apical expansion. Thus Crb3.L+ vesicles are ideally positioned for the fast delivery of a ready-made stock of membrane dedicated to apical membrane expansion.

Crb3.L depletion causes an initial reduction of the apical surface in association with decreased levels of F-actin and pERM at the apical cell aspect. MCC apical surface expansion is autonomously driven by 2D pressure powered by the apical medial F-actin pool (Sedzinski et al., 2016). Ezrin depletion in the *Xenopus* multiciliated epithelia leads to profound actin meshwork disorganization (Epting et al., 2015; Yasunaga et al., 2022). The Crb juxta membrane FERM binding domain (FBM domain) binds to the FERM domain of ERM (Médina et al., 2002; Wei et al., 2015; Whiteman et al., 2014a). In the ovarian follicle epithelium of *Drosophila*, Crb with nonfunctional FBM domain does not stabilize the activated ERM Moesin in the subapical region resulting in the loss of filamentous actin (Sherrard and Fehon, 2015). Thus, in MCCs, the most parsimonious scenario is that Crb3.L captures and stabilizes phosphorylated Ezrin in the growing apical domain, allowing progressive growth of the actin meshwork in coordination with apical membrane delivery.

### Crb3.L vesicles might regulate the contractility and dynamics of the actin meshwork in MCCs

Crumbs vesicles are the nexus of different members of the cell cytoskeleton and might bridge the rigid microtubule network to contractile acto-myosin network required for BB ascension. According to its protein partners, and the type of actin pools (medial versus junctional), Crb promotes or represses contractile activity of different acto-myosin meshworks (Biehler et al., 2021; Flores-Benitez and Knust, 2015; Ramkumar et al., 2016; Röper, 2012b; Salis et al., 2017a; Simões et al., 2022). Thus, MCCs could offer a unique model to study in the same cell how Crb could contribute to very different actin-dependent processes: apical constriction (during MCC insertion), apical expansion (during MCC emergence) and BB ascension. In *Drosophila*, Crb promotes medial actomyosin contractility during the process of salivary gland invagination in a Rab11-myosin II-dependent manner (Le and Chung, 2021). Interestingly, both Crb3.L and Rab11 accumulate in the subapical domain of MCCs at the time of their intercalation (Kim et al., 2012). In other model systems, Crb colocalizes with Rab11 endosomes (Aguilar-Aragon et al., 2020; Iioka et al., 2019; Schlüter et al., 2009). Thus, a population of Crb3.L/Rab11 doubly positive endosomes could shape the cell for optimal insertion via regulation of the internal actin meshwork contractility in a Rab11-myosin II-dependent manner.

Besides controlling contractility, Crb3.L positive vesicles might also control the dynamics and/or density of the internal actin meshwork required for BB ascension. In the mouse oocyte, the interconnected network of vesicles and actin allows the asymmetric positioning of the meiotic spindle for oocyte maturation (Holubcová et al., 2013). Rab11-positive endosomes recruit both myosin Vb and the actin nucleator Spir (Holubcová et al., 2013). Myosin V are unconventional myosins using their F-actin binding activity to transport organelles along F-actin tracks (Wong and Weisman, 2021). Hence, Rab11 vesicles are proposed to generate asymmetric forces in a myosin V-dependent manner (Holubcová et al., 2013). The number and volume of vesicles modulate the network density by clustering and sequestering the nucleators of the network, thus controlling the cortical oriented displacement of the meiotic spindle (Holubcová et al., 2013). Crb has been shown to interact with myosin V in *Drosophila*, and myosin V is recruited to Rab11 vesicles (Aguilar-Aragon et al., 2020; Dhekne et al., 2014; Lattner et al., 2019; Pocha et al., 2011; Pylypenko et al., 2016). Thus, Crb3.L vesicles accumulation in the subapical domain of the inserting MCCs could also contribute to the construction of a dynamic asymmetric actin meshwork, allowing BB ascension. In addition, in MCCs, the overexpression of a dominant negative myosin Vc construct led to BB migration defect (Tu et al., 2018). Thus, Crb3.L could control BB migration via proper targeting of a myosin V.

In conclusion, our work unveils a new aspect of Crb3 function in multiciliogenesis in vertebrates. At the tissue level, Crb3 was shown to be required for the differentiation of the proximal airway mucociliary epithelium, including MCC progenitors (Szymaniak et al., 2015). Consequently, MCCs and a fortiori cilia are absent from the proximal airway of lung targeted *crb3* knock-out mice (Szymaniak et al., 2015). This likely explains the respiratory distress syndrome of germline *crb3* knock-out mice (Charrier et al., 2015b; Szymaniak et al., 2015; Whiteman et al., 2014b). In our model, MCCs are present, but display abnormal ciliary tuft, suggesting that Crb3.L is dispensable for MCC specification in the *Xenopus* mucociliary epithelium.

At the cellular level, Crb3 appears to be involved in ciliary trafficking in MCCs as suggested by the cilia phenotype of *crb3* null zebrafish mutants (Hazime and Malicki, 2017). In our model, a ciliary trafficking defect is not excluded, but was not possible to test here as we reveal an earlier function in the emergence of the MCC apical domain in a pERM-dependent manner. Finally, this study reveals that *Xenopus* MCCs may represent a novel paradigm to understand how Crb controls different facets of actin cytoskeleton dynamics in vertebrates.

## Acknowledgements

We are grateful to E Bazellières, Grégoire Michaux, A Pasini for critical reading of the manuscript. We wish to thank PA Bidaud, A Pacquelet, J Pécréaux and JP Tassan for stimulating discussion. We thank P. Walentek for providing the CP110-GFP plasmid. We thank Xavier Pinson, Stéphanie Dutertre, Rémi Flores-Flores, Elsa Castellani for their technical assistance with confocal microscopy. We thank Florian Roguet and Julien Maurais for *Xenopus* care. We thank all members of the Gene Expression and Development team (GED), JP Tassan and Virginie Thomé for sharing *Xenopus* expertise and/or reagents.

## Authors contributions

Conceptualization: C.B, L.K, A.LB.; Methodology: C.B., L.K.; Validation: C.B, L.K A.LB; Investigation: C.B, J.R.; Writing - original draft: C.B, A.LB, L.K; Supervision: L.K.; A.LB; Project administration: A.LB; Funding acquisition: A.LB, L.K.

---

**Supplementary figure 1a:**
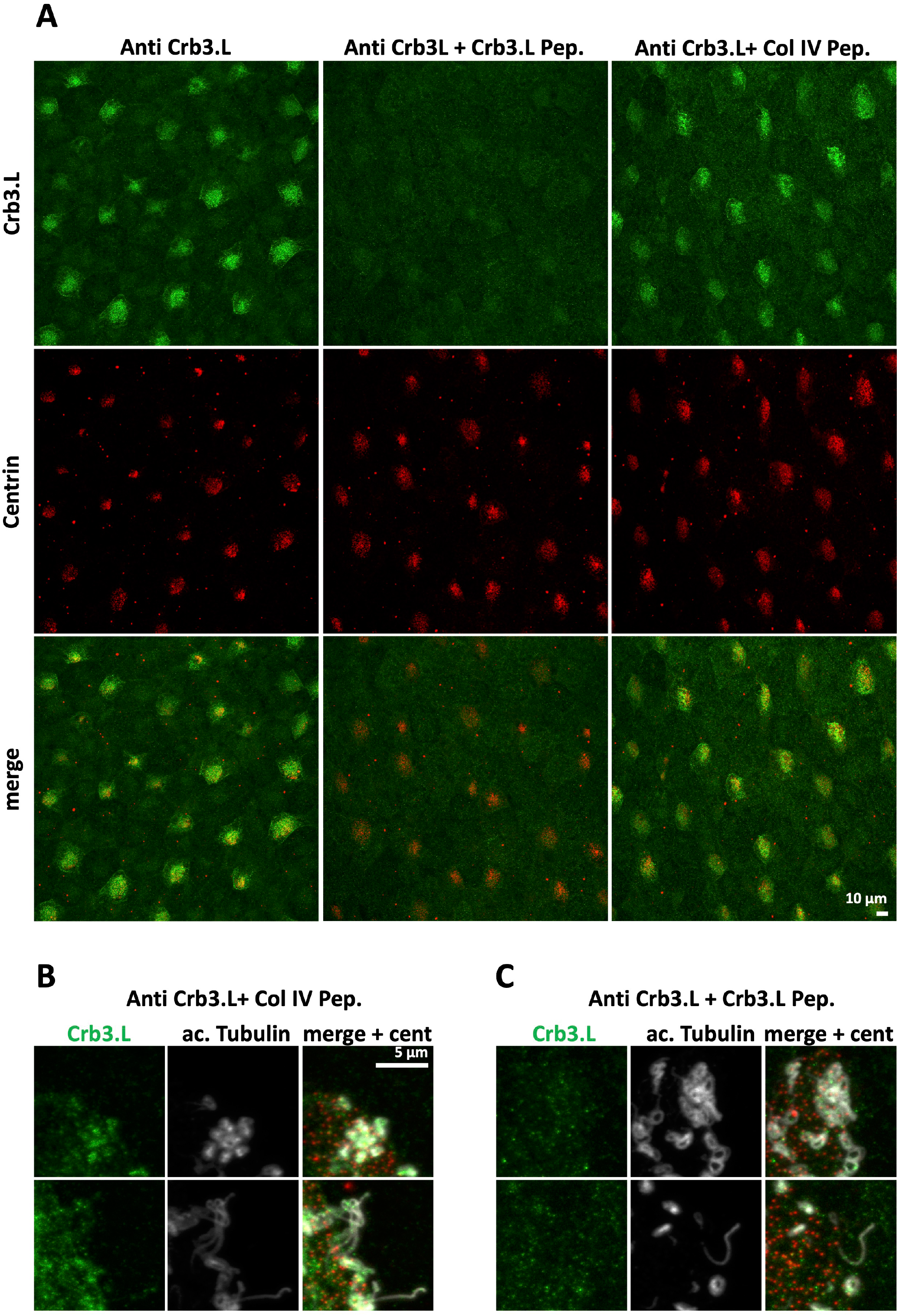
Crb3.L antibody staining is lost in immunogenic peptide competition assay. A-C: Maximal intensity projection of en face view of whole-mount St 20-21 *Xenopus laevis* embryonic mucociliary epidermis stained with anti Crb3.L antibody in control condition, in competitive condition with the addition of the immunogenic peptide, in control condition with the addition of an unrelated peptide (*Oopsacas. minuta* type IV collagen). The ratio between antibody and peptides was 1 to 25 moles. BBs were labelled with anti-Centrin antibody. Crb3.L staining in MCCs is efficiently decreased upon competition with the immunogenic peptide. B-C: Higher magnification focusing on cilia. Cilia were labelled with anti-acetylated a-Tubulin antibody. Note Crb3.L spotted staining on cilia when the Crb3.L antibody is preincubated with a none-related peptide. C: Note the disappearance of the ciliary signal upon immunogenic peptide competition.

**Supplementary figure 1b:**
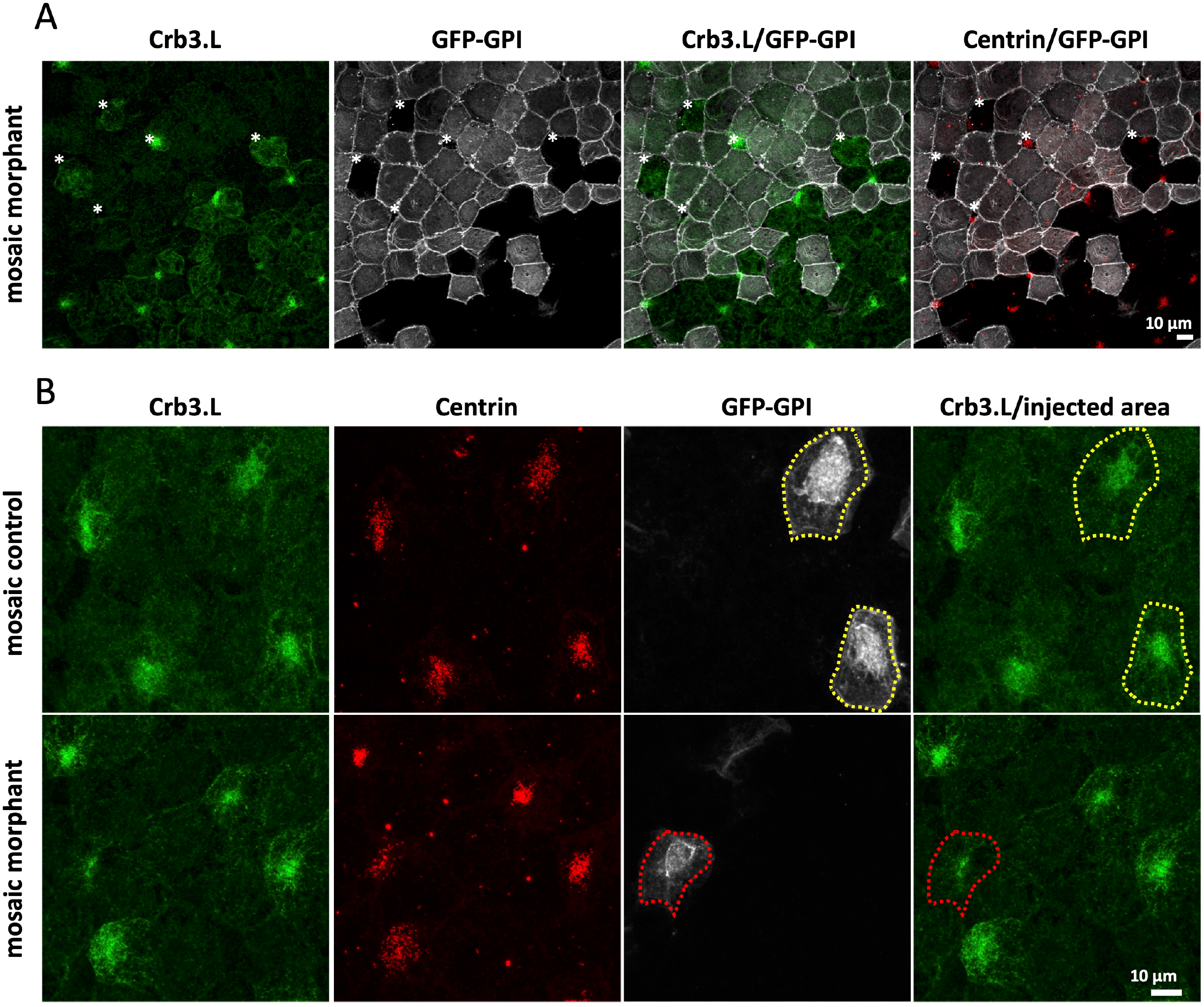
Crb3.L antibody staining decreases upon *crb3.L* morpholino depletion. A-B: Maximal intensity projection of en face view of whole-mount St 20 *Xenopus laevis* embryos. Control embryos were injected with 200 pg *GFP-GPI* mRNA to label cell membranes. Morphants were injected with *GFP-GPI* mRNA and anti *crb3.L* morpholino (15 ng of *5’UTR-crb3.L-mo*). Mosaic regions of control and morphant embryos allow easy comparison of Crb3.L staining between uninjected and tracer injected cells (in control embryo), or uninjected and Crb3.L-depleted cells (in morphants). BBs are detected with anti-Centrin antibody, cell membranes are labelled with GFP-GPI. A: mosaic morphant region showing that Crb3.L expression is globally decreased in all injected cells, including in non-MCCs cells. *indicates uninjected cells or area in otherwise *5’UTR-crb3.L-mo* injected region B: mosaic regions focusing on MCCs expanding their apical surface. Red dotted lines surround *crb3.L* morpholino and tracer injected MCCs. Yellow dotted lines surround control MCCs injected with the tracer alone.

**Supplementary figure 2:**
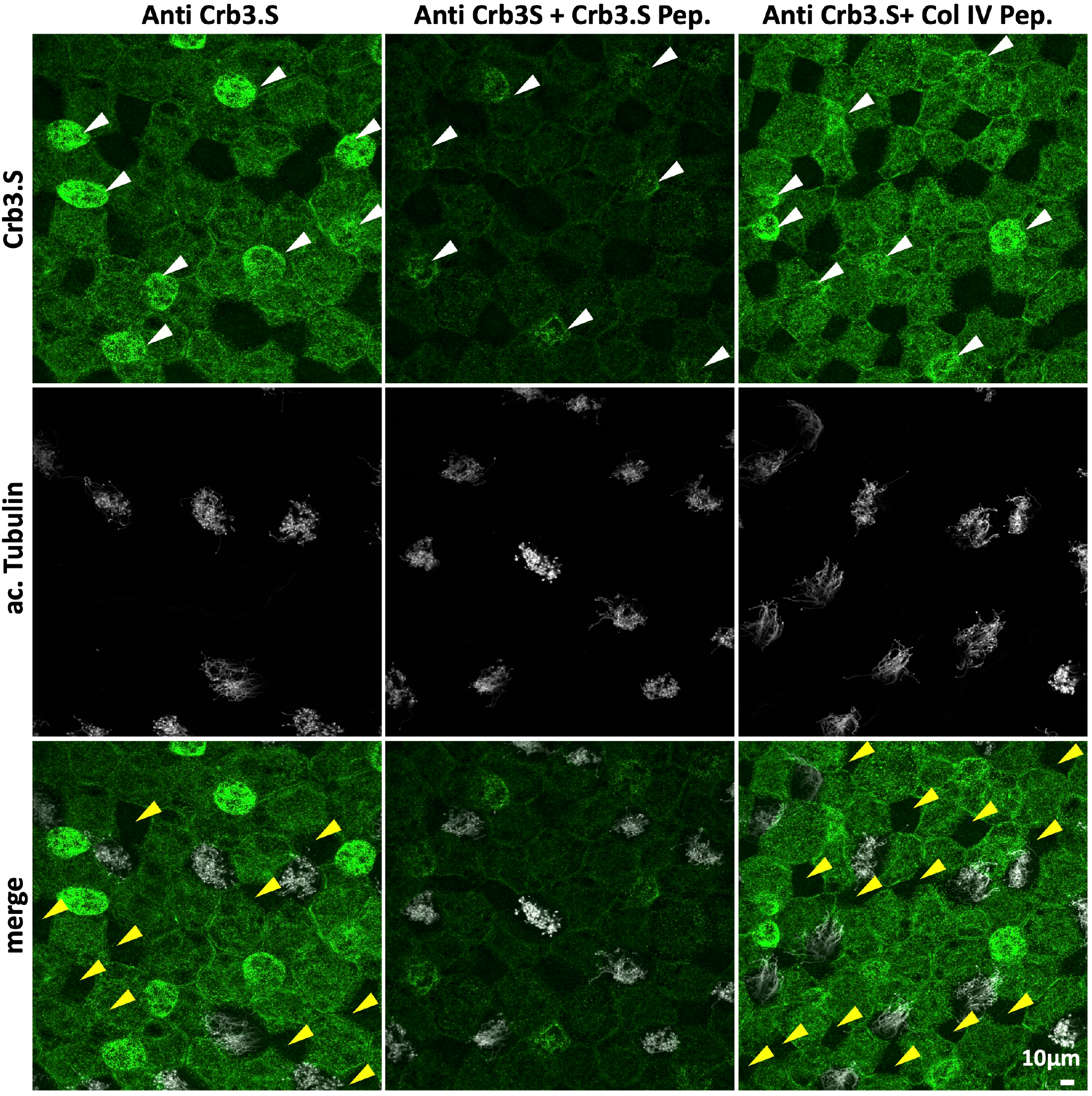
Crb3.S antibody staining decreases in immunogenic peptide competition assay. Pictures are maximal intensity projection of en face view of whole-mount St 32 heterozygous *Xenopus laevis albino* embryos stained with Crb3.S, in control condition, in competitive condition with the addition of the immunogenic peptide, in control condition with the addition of an unrelated peptide (*Oopsacas. minuta* type IV collagen). The ratio between antibody and peptides was 1 to 25 moles. White arrowheads point at SSCs, which express the highest level of Crb3.S, as revealed in Figure 2B. Yellow arrowheads point at cells of the outer layer (presumably ionocytes) with very low Crb3.S expression level. Note the drastic diminution of Crb3.S staining in the SSCs upon immunogenic peptide incubation.

**Supplementary figure 6:**
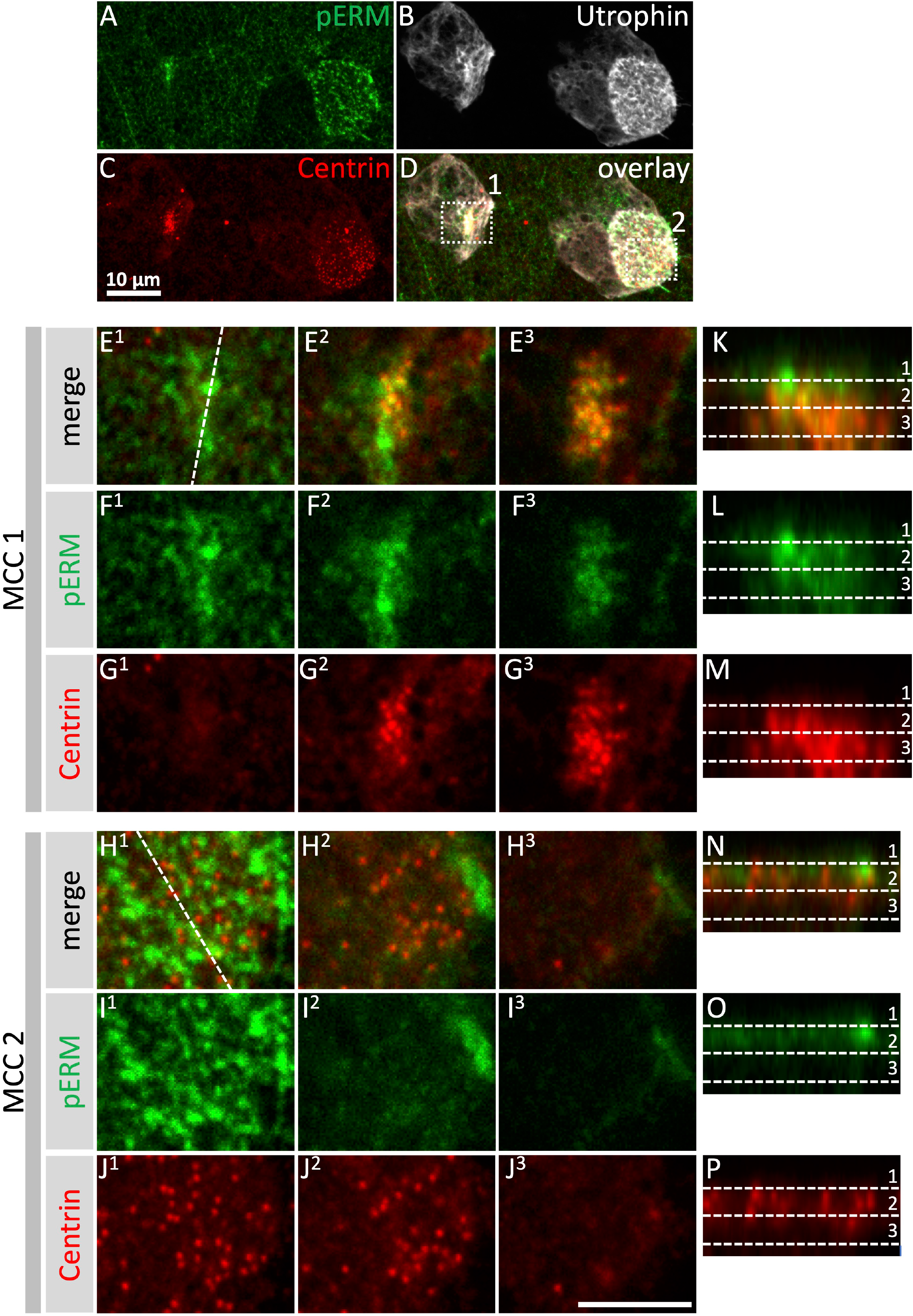
pERM is transiently recruited at BBs during their ascension. (A-D) Maximum intensity projection of early inserting MCCs in stage 18 embryos. Embryos were injected with utrophin-GFP to label actin, stained with anti-Centrin to label BBs, and anti-pERM. (D) MCC 1 is just pointing at the apical surface, MCC 2 is well engaged in apical surface expansion (135 μm^2^). The white dotted squares delimit the cropped regions of MCC 1 and MCC 2 that are used for examination of p-ERM staining at different cell levels in panels (E^1^-P). (E^1^-J^3^) confocal x, y plans of the apical surfaces and two intracytoplasmic x, y sections of increasing depth are presented for MCC 1 and MCC 2. (E^1^, F^1^, G^1^, H^1^, I^1^, J^1^) are the most apical plans. (K-P) are x, z optical confocal sections of MCC 1 and MCC 2. (E^1^-P) Precise coordinates of each confocal sections are explained below. In E^1^ the dotted line indicates the x, z axis for the optical sections presented in K, L, M. In H^1^ the dotted line indicates the x, z axis for the optical sections presented in N, O, P. The dotted lines (1, 2, 3) in (K-P) indicate the level of the x, y sections presented in (E^1^-J^3^).

